# The categorical neural organization of speech aids its perception in noise

**DOI:** 10.1101/652842

**Authors:** Gavin M. Bidelman, Lauren C. Bush, Alex M. Boudreaux

## Abstract

We investigated whether the categorical perception (CP) of speech might also provide a mechanism that aids its perception in noise. We varied signal-to-noise ratio (SNR) [clear, 0 dB, -5 dB] while listeners classified an acoustic-phonetic continuum (/u/ to /a/). Noise-related changes in behavioral categorization were only observed at the lowest SNR. Event-related brain potentials (ERPs) differentiated phonetic vs. non-phonetic (category ambiguous) speech by the P2 wave (∼180-320 ms). Paralleling behavior, neural responses to speech with clear phonetic status (i.e., continuum endpoints) were largely invariant to noise, whereas responses to ambiguous tokens declined with decreasing SNR. Results demonstrate that phonetic speech representations are more resistant to degradation than corresponding acoustic representations. Findings suggest the mere process of binning speech sounds into categories provides a robust mechanism to aid perception at the “cocktail party” by fortifying abstract categories from the acoustic signal and making the speech code more resistant to external interferences.

## 1. INTRODUCTION

A basic tenet of perceptual organization is that sensory phenomena are subject to invariance: similar features are mapped to common identities (equivalence classes) by assigning similar objects to the same membership (Goldstone & Hendrickson, 2010), a process known as categorical perception (CP). In the context of speech, CP is demonstrated when gradually morphed sounds along an equidistant acoustic continuum are heard as only a few discrete classes (Bidelman et al., 2013; Harnad, 1987; Liberman et al., 1967; Pisoni, 1973; Pisoni & Luce, 1987). Equal physical steps along a signal dimension do not produce equivalent changes in percept (Holt & Lotto, 2006). Rather, listeners treat sounds within a given category as perceptually similar despite their otherwise dissimilar acoustics. Skilled categorization is particularly important for spoken and written language, as evidenced by its role in reading acquisition (Mody et al., 1997; Werker & Tees, 1987), sound-to-meaning learning (Myers & Swan, 2012; Reetzke et al., 2018), and putative deficits in language-based learning disorders (e.g., specific language impariment, dyslexia; Calcus et al., 2016; Noordenbos & Serniclaes, 2015; Werker & Tees, 1987). To arrive at categorical decisions, acoustic cues are presumably weighted and compared against internalized “templates” in the brain, built through repetitive exposure to one’s native language (Bidelman & Lee, 2015; Guenther et al., 2004; Iverson et al., 2003a; Kuhl, 1991).

Beyond providing observers a smaller, more manageable perceptual space, CP might also aid degraded speech perception if phonetic categories are somehow more resistant to noise (Gifford et al., 2014; Helie, 2017). Indeed, categories (a higher-level code) are thought to be more robust to noise degradations than physical surface features of a signal (lower-level sensory code) (Bidelman et al., under review; Helie, 2017). A theoretical example of how categorical processing might aid the perception of degraded speech is illustrated in Figure 1.

**Figure 1:**
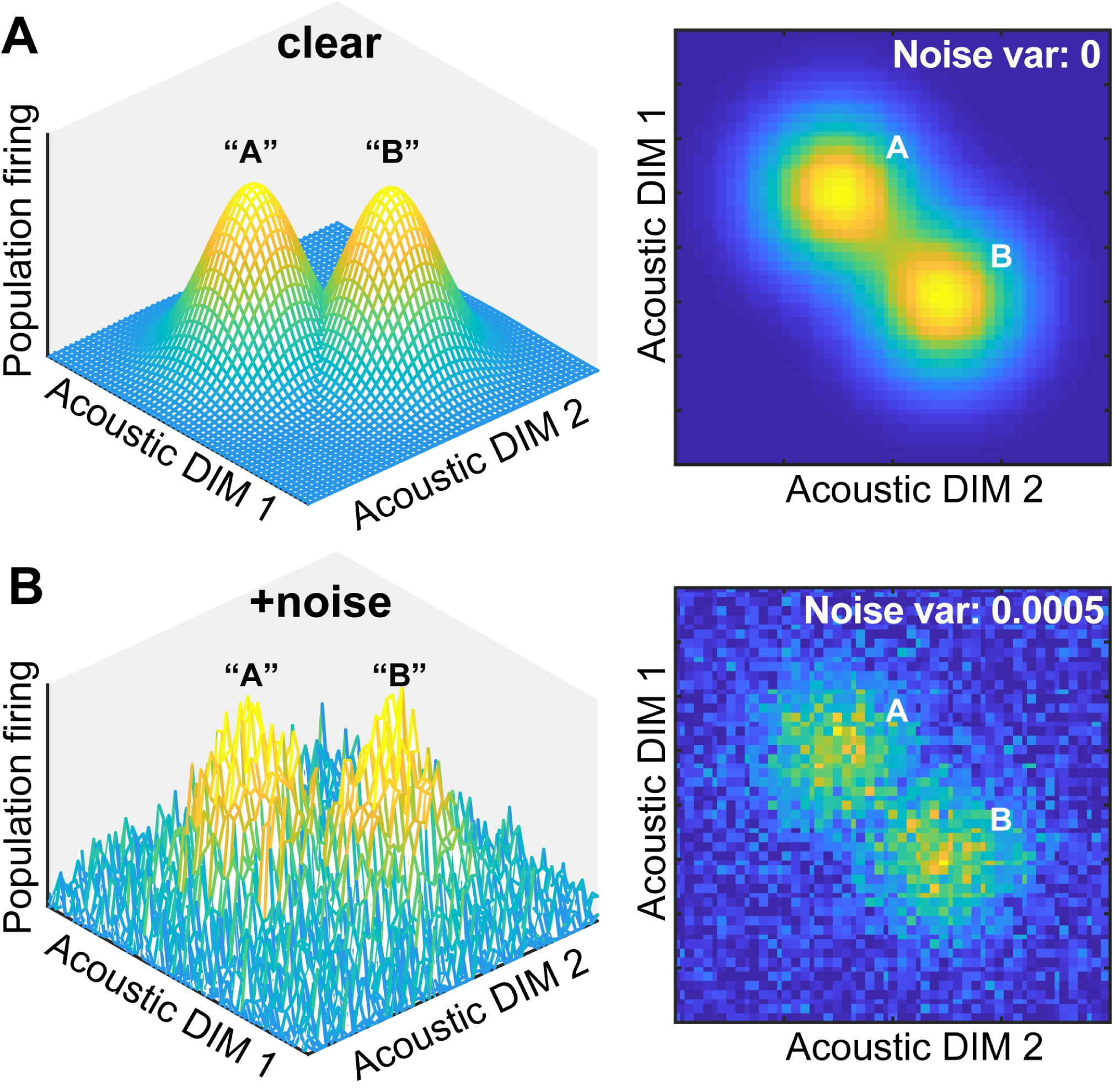
Theoretical framework for noise-related influences on categorical speech representations. (**A**) The neural representation of speech is modeled as a multidimensional feature space where populations of auditory cortical neurons code different dimensions (DIM) of the input. DIMS here are arbitrary but could reflect any behaviorally relevant feature of speech (e.g., F0, duration, etc.) Both 3D and 2D representations are depicted here for two stimulus classes. Categorical coding (modeled as a Gaussian mixture) is reflected by an increase in local firing rate for perceptually similar stimuli (“A” and “B”). (**B**) Noise blurs physical acoustic details yet spares categories as evidenced by the resilience of the peaks in neural space. Neural noise was modeled by changing the variance of additive Gaussian white noise.

Consider the neural representation of speech as a multidimensional feature space. Populations of auditory cortical neurons code different dimensions of the acoustic input. Categorical coding could be reflected as an increase (or conversely, decrease) in local firing rate for stimuli that are perceptually similar despite their otherwise dissimilar acoustics (“A” and “B”) (e.g., Guenther & Gjaja, 1996; Guenther et al., 2004; Recanzone et al., 1993). Although noise interference would blur physical acoustic details and create a noisier cortical map, categories would be partially spared—indicated by the remaining “peakedness” in the neural space. Thus, both the construction of perceptual objects and natural discrete binning process of CP might enable category members to “pop out” among a noisy feature space (e.g., Nothdurft, 1991; Perez-Gay et al., 2018). Consequently, the mere process of grouping speech sounds into categories might aid comprehension of speech-in-noise (SIN)—assuming those representations are not too severely compromised and remain distinguishable from noise itself. This theoretical framework provides the basis for the current empirical study.

Building on our recent efforts to decipher the neurobiology of “cocktail party” listening and understand the physiological mechanisms supporting robust speech perception (for review, see Bidelman, 2017), this study aimed to test whether speech sounds carrying strong phonetic categories are more resilient to the deleterious effects of noise than categorically ambiguous speech sounds. When category-relevant dimensions are less distinct and perceptual boundaries are particularly noisy, additional mechanisms for enhancing separation must be engaged (Livingston et al., 1998). We hypothesized the phonetic groupings inherent to speech may be one such mechanism. Because phonetic categories reflect a more abstract, higher-level representation of speech (i.e., acoustic + phonetic code), we reasoned they would be more robust to noise than physical features of speech that do not engage phonetic-level processing (i.e., acoustic code) (cf. Bidelman et al., under review; Helie, 2017). To test this possibility, we recorded high-density event-related potentials (ERPs) while listeners categorized speech continua in different levels of acoustic noise. The critical comparison was between responses to stimuli at the endpoints vs. midpoint of the acoustic-phonetic continuum. We predicted that if the categorization process aids figure-ground perception, speech tokens having a clear phonetic identity (continuum endpoints) would elicit lesser noise-related change in the ERPs than non-phonetic tokens (continuum midpoint), which have a bistable (ambiguous) percept and lack a clear phonetic identity.

## 2. MATERIALS & METHODS

### 2.1 Participants

Fifteen young adults (3 male, 12 females; age: *M* = 24.3, *SD* = 1.7 years) were recruited from the University of Memphis student body. All exhibited normal hearing sensitivity confirmed via a threshold screening (i.e., < 20 dB HL, audiometric frequencies 250 - 8000 Hz). Each participant was strongly right-handed (87.0± 18.2% laterality index; Oldfield, 1971) and had obtained a collegiate level of education (17.8±1.9 years). Musical training is known to modulate categorical processing and SIN listening abilities (Bidelman & Alain, 2015b; Bidelman & Krishnan, 2010; Bidelman et al., 2014b; Parbery-Clark et al., 2009; Yoo & Bidelman, under review). Consequently, we required that all participants had minimal music training throughout their lifetime (mean years of training: 1.3±1.8 years). All were paid for their time and gave informed consent in compliance with the Declaration of Helsinki and a protocol approved by the Institutional Review Board at the University of Memphis.

### 2.2 Speech continuum and behavioral task

We used a synthetic five-step vowel continuum to assess the neural correlates of CP (Bidelman & Alain, 2015b; Bidelman & Walker, 2017; Bidelman et al., 2014b). Each token of the continuum was separated by equidistant steps acoustically based on first formant frequency (F1), but was perceived categorically from /u/ to /a/. Tokens were 100 ms, including 10 ms of rise/fall time to reduce spectral splatter in the stimuli. Each contained identical voice fundamental (F0), second (F2), and third formant (F3) frequencies (F0: 150, F2: 1090, and F3: 2350 Hz). The F1 was parameterized over five equal steps between 430 and 730 Hz such that the resultant stimulus set spanned a perceptual phonetic continuum from /u/ to /a/ (Bidelman et al., 2013). Speech stimuli were delivered binaurally at 75 dB SPL through shielded insert earphones (ER-2; Etymotic Research) coupled to a TDT RP2 processor (Tucker Davis Technologies).

This same speech continuum was presented in one of three noise blocks varying in signal-to-noise ratio (SNR): clear, 0 dB SNR, -5 dB SNR (Fig. 2). These noise levels were selected based on extensive pilot testing which confirmed they differentially hindered speech perception. The masker was a speech-shaped noise based on the long-term power spectrum of the vowel set. Noise was presented continuously so it was not time-locked to the stimulus presentation, providing a constant backdrop of acoustic interference during the categorization task (e.g., Alain et al., 2012; Bidelman et al., 2018; Bidelman & Howell, 2016). SNR was manipulated by changing the level of the masker to ensure SNR was inversely correlated with overall sound level (Binder et al., 2004). Noise block order was randomized within and between participants.

**Figure 2:**
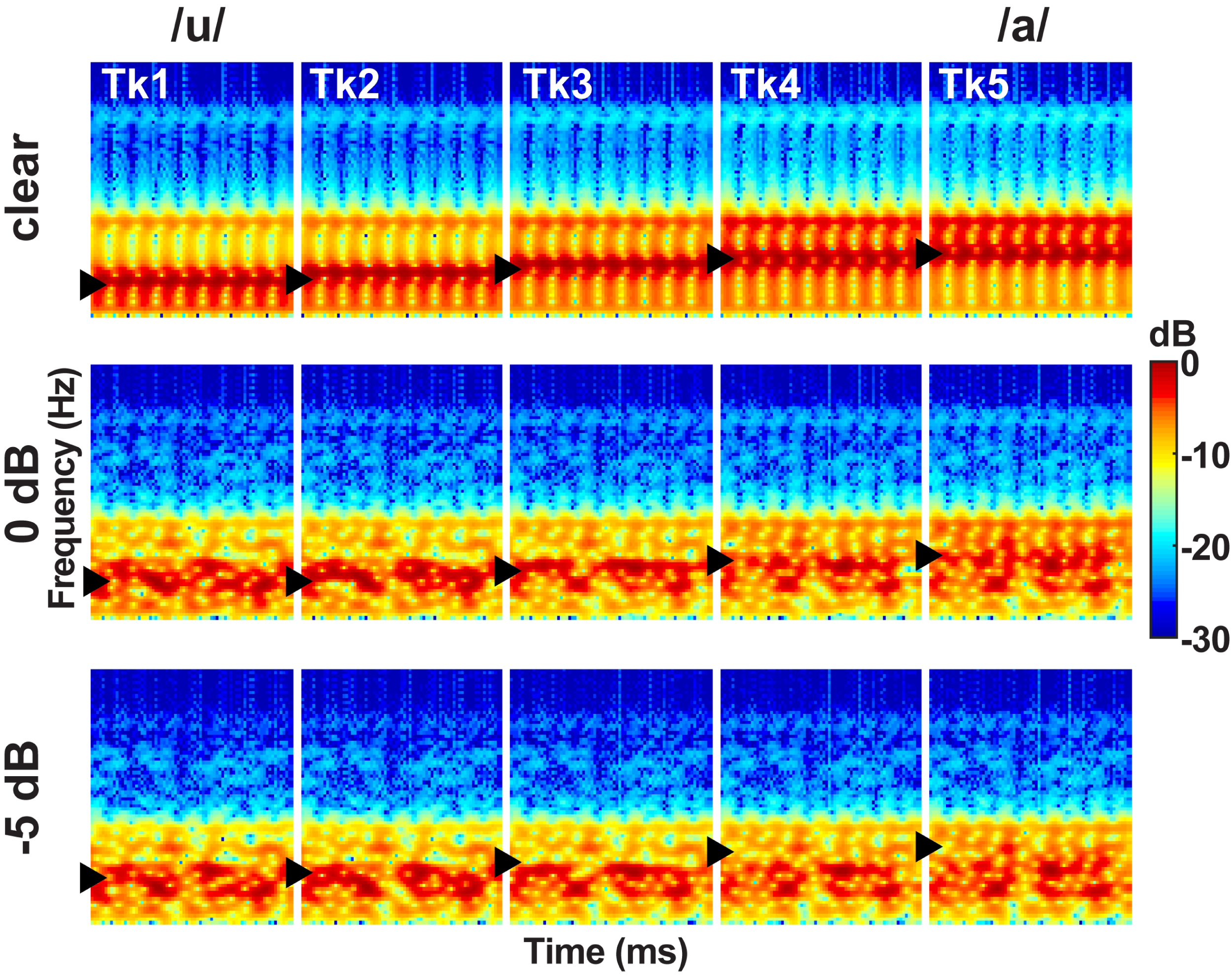
Acoustic spectrograms of the speech continuum as a function of SNR. Vowel first formant frequency was parameterized over five equal steps (430 to 730 Hz, ►), resulting in a perceptual phonetic continuum from /u/ to /a/. Token durations were 100 ms. Speech stimuli were presented at 75 dB SPL with noise added parametrically to vary SNR.

The task was otherwise identical to our previous neuroimaging studies on CP (e.g., Bidelman & Alain, 2015b; Bidelman et al., 2013; Bidelman & Walker, 2017). During EEG recording, listeners heard 150 trials of each individual speech token (per noise block). On each trial, they were asked to label the sound with a binary response (“u” or “a”) as quickly and accurately as possible. Following the listener’s behavioral response, the interstimulus interval (ISI) was jittered randomly between 800 and 1000 ms (20 ms steps, uniform distribution) to avoid rhythmic entrainment of the EEG and the anticipation of subsequent stimuli.

### 2.3 EEG recording and preprocessing

EEGs were recorded from 64 sintered Ag/AgCl electrodes at standard 10-10 scalp locations (Oostenveld & Praamstra, 2001). Continuous data were digitized using a sampling rate of 500 Hz (SynAmps RT amplifiers; Compumedics Neuroscan) and an online passband of DC-200 Hz. Electrodes placed on the outer canthi of the eyes and the superior and inferior orbit monitored ocular movements. Contact impedances were maintained < 10 kΩ during data collection. During acquisition, electrodes were referenced to an additional sensor placed ∼ 1 cm posterior to the Cz channel. EEG pre-processing was performed in BESA® Research (v7) (BESA, GmbH). Ocular artifacts (saccades and blinks) were first corrected in the continuous EEG using a principal component analysis (PCA) (Picton et al., 2000). Cleaned EEGs were then filtered (1-30 Hz), epoched (-200-800 ms), baseline corrected to the pre-stimulus interval, and averaged in the time domain resulting in 15 ERP waveforms per participant (5 tokens * 3 noise conditions). For analysis, data were re-referenced using BESA’s reference-free virtual montage. This montage computes a spherical spline-interpolated voltage (Perrin et al., 1989) for each channel relative to the mean voltage over 642 equidistant locations covering the entire sphere of the head. This montage is akin to common average referencing but results in a closer approximation to true reference free waveforms (Scherg et al., 2002). However, reported results were similar using a common average reference (data not shown).

ERP quantification focused on the latency range of the P2 wave as previous studies have shown the neural correlates of CP emerge around the timeframe of this component (Bidelman & Alain, 2015b; Bidelman & Lee, 2015; Bidelman et al., 2013; Bidelman & Walker, 2017). Guided by visual inspection of grand averaged data, we measured the amplitude of the evoked potentials as the positive-going deflection between 180-320 ms. This window covered both the P2 and following P3b-like deflections (see Fig. 4). To evaluate whether ERPs showed categorical coding, we averaged response amplitudes to prototypical tokens at the endpoints of the continuum and compared this combination to the ambiguous token at its midpoint (e.g., Bidelman, 2015; Bidelman & Walker, 2017; Liebenthal et al., 2010). This contrast (i.e., mean[Tk1, Tk5] vs. Tk3) allowed us to assess the degree to which noise affected the neural encoding of phonetic categories (Tk1/5) vs. non-phonetic (Tk3) speech sounds and thus affected categorical processing. The rationale for this analysis is that it effectively minimizes stimulus-related differences in the ERPs, thereby isolating categorical/perceptual processing. For example, Tk1 and Tk5 are expected to produce distinct ERPs due to exogenous processing alone. However, comparing the average of these responses (i.e., mean[Tk1, Tk5]) to that of Tk3 allows us to better isolate ERP modulations related to categorical coding (Bidelman & Walker, 2017).

Averaging endpoint responses doubles the number of trials for the prototypes relative to the ambiguous condition, which could mean differences were attributable to SNR of the ERPs rather than CP effects, *per se* (Hu et al., 2010). To rule out this possibility, we measured the SNR of the ERPs as 10log(RMS_ERP_/RMS_baseline_) (Bidelman, 2018) where RMS_ERP_ and RMS_baseline_ were the RMS amplitudes of the ERP (signal) portion of the epoch window (0-800 ms) and pre-response baseline period (-200-0ms ms), respectively. Critically, SNR of the ERPs did not differ across conditions [*F_5,70_* =0.56, *p*=0.73], indicating that neural activity was not inherently noisier for a given token type or acoustic noise level.

### 2.4 Behavioral data analysis

Identification scores were fit with a sigmoid function *P* = 1/[1+*e*^-*β1*(*x* - *β0*)^], where *P* is the proportion of trials identified as a given vowel, *x* is the step number along the stimulus continuum, and *β_0_* and *β_1_* the location and slope of the logistic fit estimated using nonlinear least-squares regression. Comparing parameters between SNR conditions revealed possible differences in the location and “steepness” (i.e., rate of change) of the categorical boundary as a function of noise degradation. Larger *β_1_* values reflect steeper psychometric functions and thus stronger categorical perception.

Behavioral speech labeling speeds (i.e., reaction times [RTs]) were computed as listeners’ median response latency across trials for a given condition. RTs outside 250-2500 ms were deemed outliers (e.g., fast guesses, lapses of attention) and were excluded from the analysis (Bidelman et al., 2013; Bidelman & Walker, 2017).

### 2.5 Statistical analysis

Unless otherwise noted, dependent measures were analyzed using a one-way, mixed model ANOVA (subject=random factor) with fixed effects of SNR (3 levels: clear, 0 dB, -5dB) and token [5 levels: Tk1-5] (PROC GLIMMIX, SAS® 9.4; SAS Institute, Inc.). Tukey-Kramer adjustments controlled Type I error inflation for multiple comparisons. The α-level for significance was *p*=0.05. We used robust linear regression as implemented by the ‘fitlm’ function in MATLAB to assess links between neural and behavioral measures.

## 3. RESULTS

### 3.1 Behavioral identification (%, RTs)

Behavioral identification functions are shown across the different noise SNRs in Figure 3A. Listeners’ identification was more categorical (i.e., dichotomous) for clear speech and became more continuous with poorer SNR. Analysis of the slopes (*β_1_*) confirmed a main effect of SNR [*F_2,28_*=35.25, *p*<0.0001] (Fig. 3B). Tukey-Kramer contrasts revealed psychometric slopes were unaltered for 0 dB SNR relative to clear speech (*p*=0.33). However, -5 dB SNR noise weakened categorization, flattening the psychometric function (-5dB vs. 0 dB, *p*<0.0001). These findings indicate the strength of categorical representations is resistant to acoustic interference. That is, even when signal and noise compete at equivalent levels, categorical processing persists. CP is weakened only for severely degraded speech (i.e., negative SNRs) where the noise exceeds the target signal.

**Figure 3:**
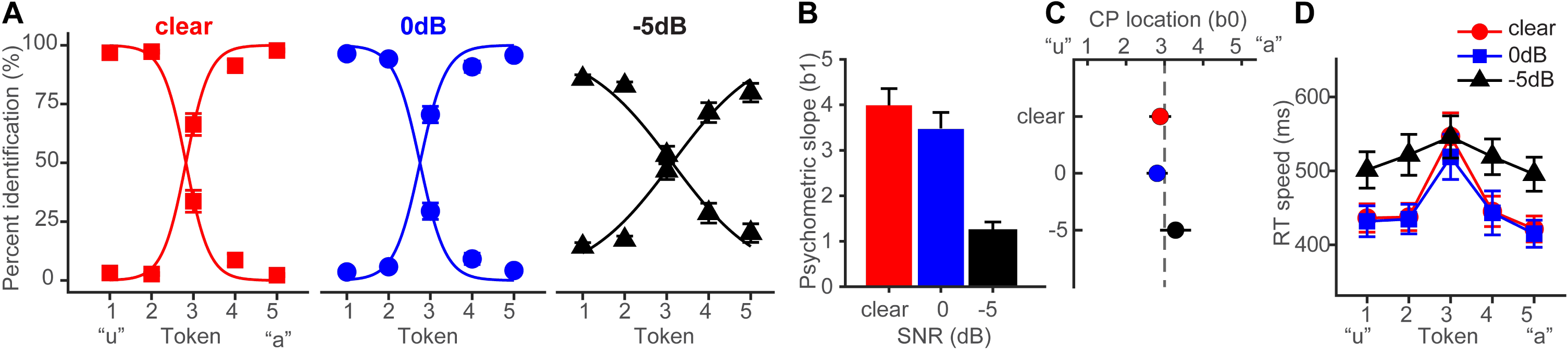
Behavioral speech categorization is robust to noise interference. (**A**) Perceptual psychometric functions for clear and degraded speech identification. Curves show an abrupt shift in perception when classifying speech indicative of discrete perception (i.e., CP). (**B**) Slopes and (**C**) locations of the perceptual boundary show speech categorization is robust even down to 0 dB SNR; the strength of CP diminishes only for highly impoverished signals as indicated by the shallow slope and slight rightward bias of the curves (i.e., more frequently responding “u”) in the -5 dB SNR condition. (**D**) Speech classification speeds (RTs) show a categorical pattern for clear and 0 dB SNR speech; participants are slower at labeling ambiguous tokens (midpoint) relative to those with a clear phonetic label (endpoints) (Bidelman & Walker, 2017; Pisoni & Tash, 1974). A categorical RT effect is not observed for highly degraded speech (-5 dB SNR). errorbars = ± s.e.m.

Noise-related changes in the psychometric function could be related to uncertainty in category distributions (prior probabilities) (Gifford et al., 2014) or lapses of attention due to task difficulty rather than a weakening of speech categories, *per se* (Bidelman et al., under review). To rule out this latter possibility, we used Bayesian inference (psignifit toolbox; Schütt et al., 2016) to estimate individual lapse (λ) and guess (γ) rates from participants’ identification data. Lapse rate (λ) was computed as the difference between the upper asymptote of the psychometric function and 100%, reflecting the probability of an “incorrect” response at infinitely high stimulus levels (i.e., responding “u” for Tk5; see Fig. 3A). Guess rate (γ) was defined as the difference between the lower asymptote and 0. For an ideal observer λ=0 and γ=0. We found neither lapse [*F_2,28_* =2.41, *p*=0.11] nor guess rate [*F_2,28_* =1.45, *p*=0.25] were modulated by SNR. This helps confirm that while (severe) noise weakened CP for speech (Fig. 3B), those effects were not driven by a lack of task vigilance or guessing, *per se* (Bidelman et al., under review; Schütt et al., 2016).

The location of the perceptual boundary (Fig. 3C) varied marginally with SNR but the shift was significant [*F_2,28_* =5.62, *p*=0.0089]. Relative to the clear condition, -5 dB SNR speech shifted the perceptual boundary rightward (*p*=0.011). This indicates a small but measurable bias to report “u” (i.e., more frequent Tk1-2 responses) in the noisiest listening condition.

Behavioral RTs, reflecting the speed of categorization, are shown in Figure 3D. An ANOVA revealed RTs were modulated by both SNR [*F_2,200_*=11.90, *p*<0.0001] and token [*F_4,200_* =5.36, *p*=0.0004]. RTs were similar when classifying clear and 0 dB SNR speech (*p*=1.0) but slowed in the -5 dB condition (*p*<0.0001). Notably, *a priori* contrasts revealed this noise-related slowing in RTs was most prominent at the phonetic endpoints of the continuum (Tk1-2 and Tk4-5); at the ambiguous (non-phonetic) Tk3, RTs were identical across SNRs (*p*s > 0.69). This suggests that the observed RT effects in noise are probably not due to a general slowing of decision speed (e.g., attentional lapses) across the board but rather, are restricted to accessing categorical representations.

CP is also characterized by a slowing in RTs near the ambiguous midpoint of the continuum (Bidelman et al., 2013; Bidelman & Walker, 2017; Bidelman et al., 2014b; Pisoni & Tash, 1974; Poeppel et al., 2004; Reetzke et al., 2018). Planned contrasts revealed this characteristic slowing in RTs for the clear [mean(Tk1,2,4,5) vs. Tk3; *p*=0.0003] and 0 dB SNR (*p*=0.0061) conditions. This categorical RT pattern was not observed at -5 dB SNR (*p*=0.59). Collectively, our behavioral results suggest noise weakened the strength of CP in both the quality and speed of categorical decisions but only when speech was severely degraded. Perceptual access to categories was otherwise unaffected by low-level noise (i.e., ≥ 0 dB SNR).

### 3.2 Electrophysiological data

Grand average ERPs are shown across tokens and SNRs in Figure 4. ERPs showed the most SNR- and token-related modulations starting around the P2 wave (∼180 ms) that persisted for another 200 ms. Visual inspection of the data indicated these modulations were most prominent at centro-parietal scalp locations. The enhanced positivity at these electrode sites following the auditory P2 might partly reflect differences in P3b amplitude (Alain et al., 2001). To quantify these effects, we measured the mean amplitudes in the 180-320 ms time window at the vertex channel (Cz) (Fig. 5). To assess the degree to which ERPs showed categorical-level coding, we then pooled tokens Tk1 and Tk5 (those with clear phonetic identities) and compared these responses to the ambiguous Tk3 at the midpoint of the continuum (Bidelman, 2015; Bidelman & Walker, 2017). An ANOVA conducted on ERP amplitudes showed responses were strongly modulated by SNR [*F_2,70_*=8.54, *p*=0.0005] and whether not the stimulus carried a phonetic label [Tk1/5 vs. Tk3: *F_1,70_* =19.11, *p* <0.0001] (Fig. 5B). Planned contrasts by SNR revealed that neural activity differentiated phonetic vs. non-phonetic speech at clear (*p*=0.0170) and 0 dB (*p*=0.0011) SNRs, but not at -5 dB (*p*=0.0915). Across SNRs, ERPs to phonetic tokens were largely resilient to noise [Tk1/5; linear contrast of SNR: *t_7_*_0_ = -2.17, *p*=0.07)]. In contrast, responses declined for non-phonetic speech sounds [Tk3; *t_7_*_0_ = -2.91, *p*=0.0098]. This indicates that that noise differentially affected the representation of category prototypes relative to phonetically ambiguous speech. Pooled across speech tokens, CLARA source analysis (Alain et al., 2017; Bidelman et al., 2018; Iordanov et al., 2014) localized sensor-level activity to bilateral sources in anterior superior temporal gyrus (STG) (cf. Bidelman & Lee, 2015). These neural findings parallel our behavioral results and suggest the categorical (phonetic) representations of speech (near STG) are more resistant to noise than those that do not carry a clear linguistic-phonetic identity.

**Figure 4:**
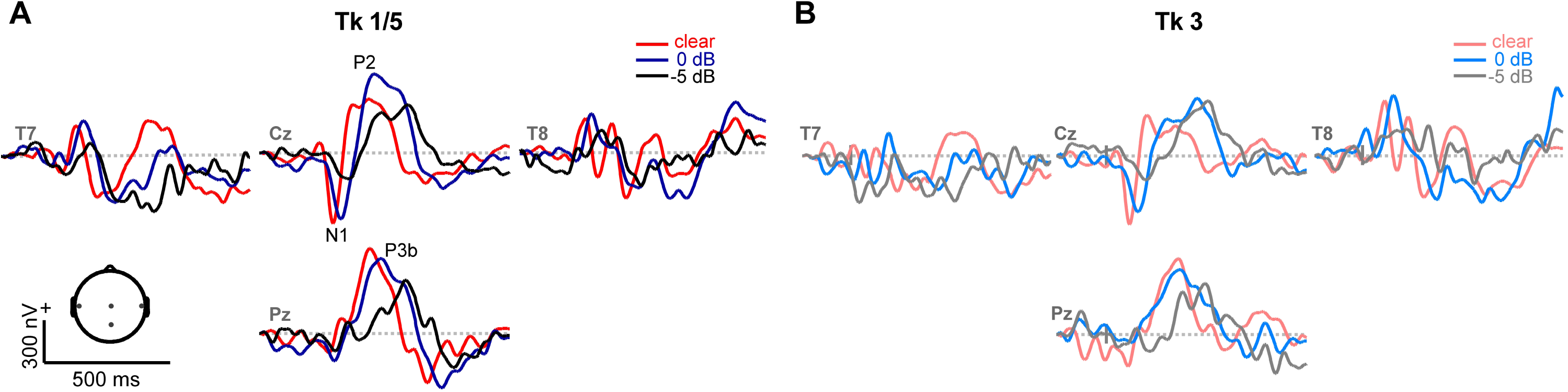
Speech-ERPs as a function of speech token and noise (SNR). Representative electrodes at central (Cz), temporal (T7/8) and parietal (Pz) scalp sites. Stimulus and noise-related modulations are most prominent between P2 and P3b (180-320 ms). (**A)** Phonetic speech tokens (Tk1, Tk5) elicit stronger ERPs than **(B)** ambiguous sounds without a clear category (Tk3). Noise weakens and prolongs the neural encoding of speech.

**Figure 5:**
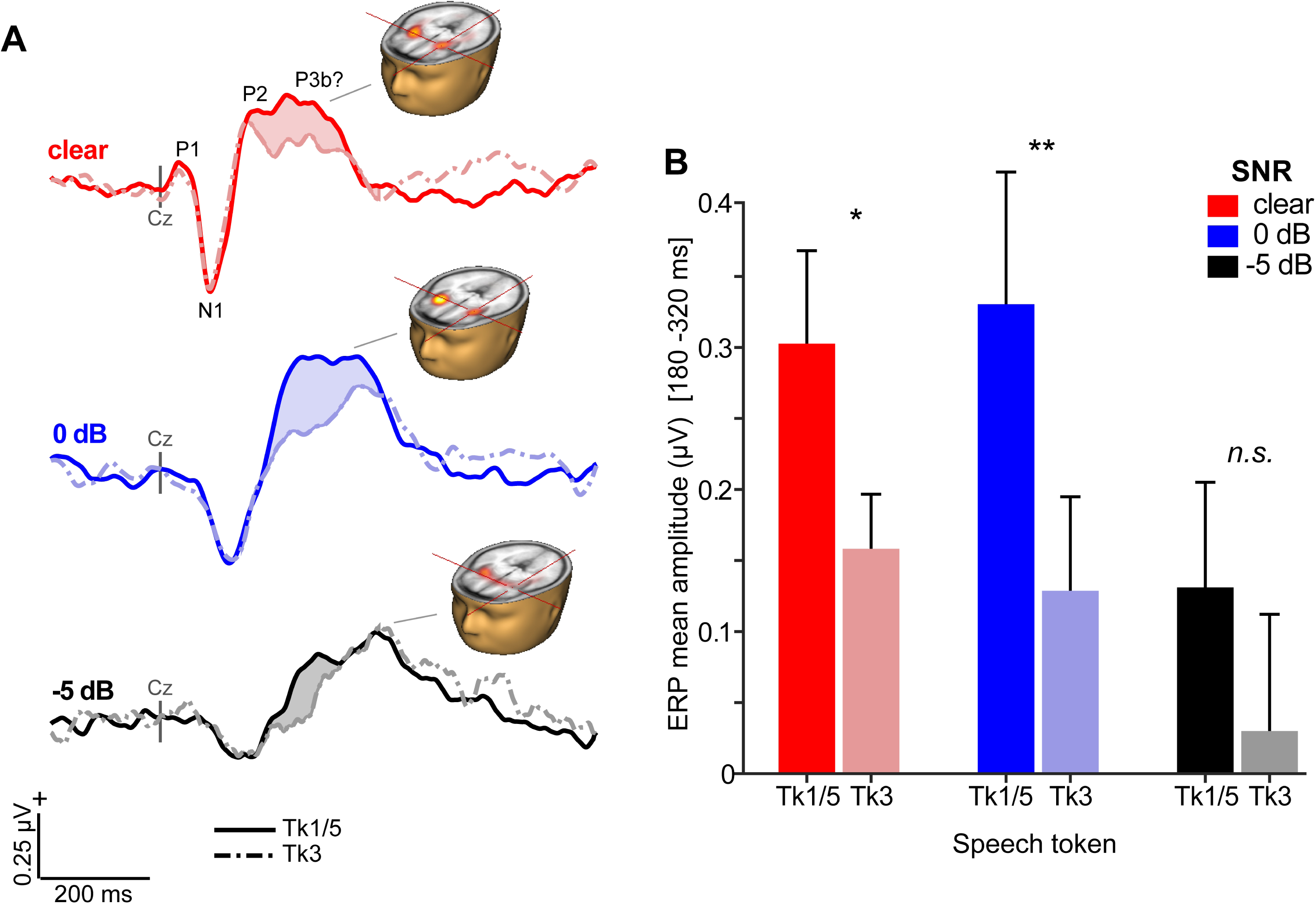
Categorical neural organization limits the degradative effects of noise on cortical speech processing. (**A**) Scalp auditory ERP waveforms (Cz electrode). (*shaded regions*) Stronger responses are observed for phonetic exemplar vs. ambiguous speech tokens [i.e., mean(Tk1, Tk5) > Tk3] but this effect varies with SNR. Brain volumes show Cortical Low resolution electromagnetic tomography Analysis Recursively Applied (CLARA; BESA® v7) (Iordanov et al., 2014) functional maps projected onto the BESA template brain (Richards et al., 2016) [latencies: 240 ms (clear), 272 ms (0 dB), 300 ms (-5 dB)]. CLARA activations (pooled across all speech tokens) localize ERP activity to anterior superior temporal cortex bilaterally. As in sensor (scalp) responses (cf. Fig. 4), source activations diminish with increasing noise (i.e., lower SNRs). (**B**) Mean ERP amplitude (180-320 ms window) is modulated by SNR and phonetic status. Categorical neural encoding (Tk1/5 > Tk3) is observed for all but the noisiest listening condition. errorbars = ± s.e.m. \**p*<0.05; ** *p*<0.01.

### 3.3 Brain-behavior relationships

The effects of noise on categorical neural processing closely paralleled the perceptual data. Figure 6A shows the group mean performance on the behavioral identification task and group mean ERP amplitudes (180-320 ms window) to the phonetic speech tokens (Tk1/5). For ease of comparison, both the neural and behavioral measures were normalized for each participant (Alain et al., 2001), with 1.0 reflecting the largest displacement in ERP amplitude and psychometric slopes, respectively. The remarkably similar pattern between brain and behavioral data imply that perceptual identification performance is predicted by the underlying neural representations for speech, as reflected in the ERPs. Indeed, correlational analyses revealed a strong association between behavioral responses and ERPs elicited by the phonetic (Tk1/5) (Fig. 6B; *R^2^* = 0.15, *p*=0.0085) but not ambiguous (Tk3) tokens (Fig. 6C; *R^2^* = 0.03, *p*=0.23). That is, more robust neural activity predicted steeper psychometric functions. These findings suggest the neural processing of speech sounds carrying clear phonetic labels predicts more dichotomous categorical decisions at the behavioral level; whereas neural responses to ambiguous (non-categorical/non-phonetic) speech tokens do not predict perceptual categorization.

**Figure 6:**
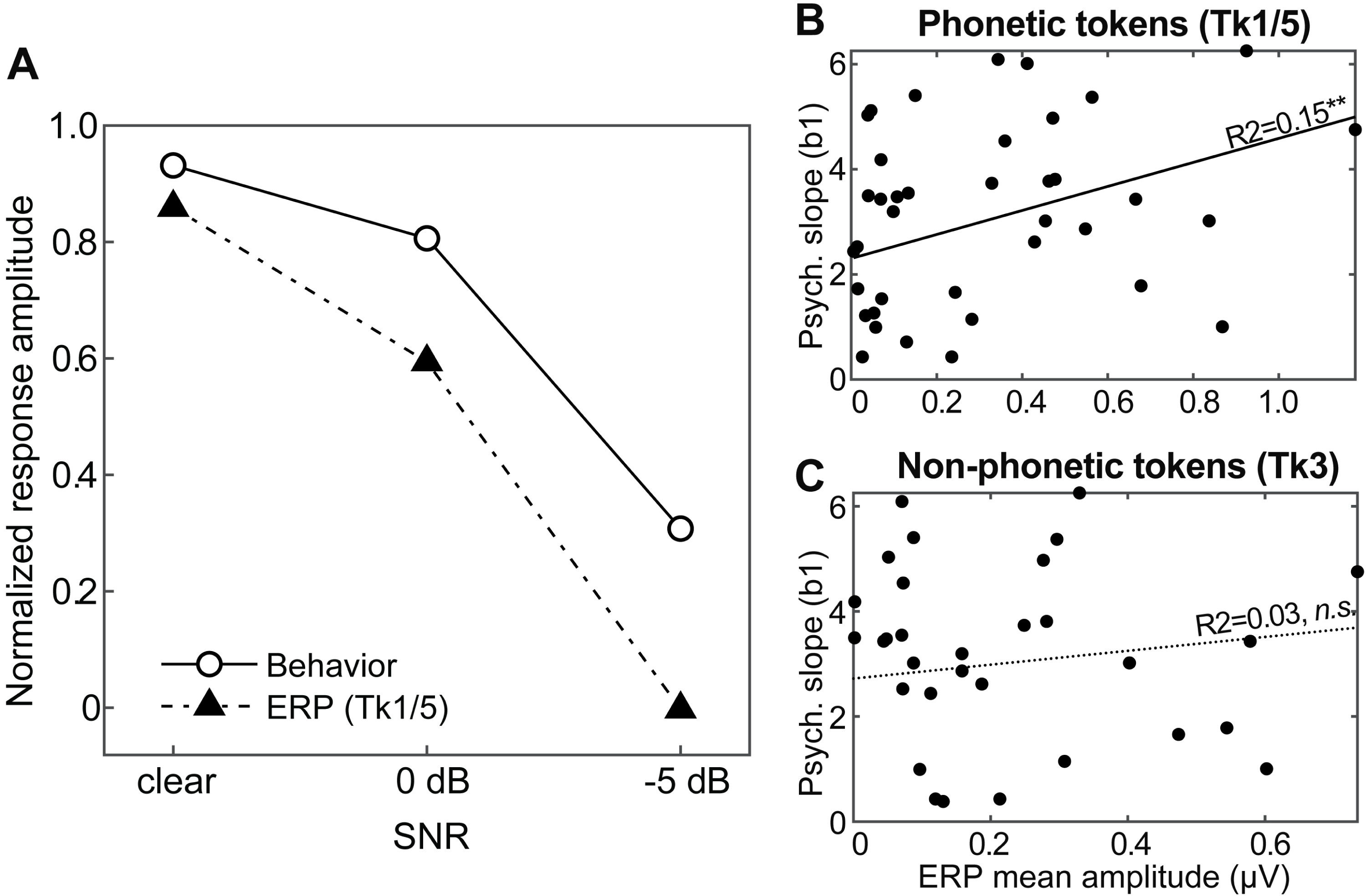
Brain-behavior associations in categorical speech perception. (**A**) Amplitudes of the auditory ERPs overlaid with behavioral data (psychometric slopes). Neural and behavioral measures are normalized for each participant (Alain et al., 2001), with 1.0 reflecting the largest displacement in ERP amplitude (180-320 ms) and psychometric slopes, respectively. (**B-C**) Correlations between behavioral CP and neural responses to (**B**) phonetic (Tk1/5) and (**C**) non-phonetic speech tokens (Tk3). Data are aggregated across SNRs (n=45 observations). Behavioral CP is predicted only by neural activity to phonetic tokens; larger ERP amplitudes elicited by Tk1/5 speech are associated with steeper, more dichotomous CP. Solid lines, significant regression; Dotted lines, non-significant regression. ** *p*<0.01.

## 4. DISCUSSION

By measuring neuroelectric activity during rapid classification of SIN, our results reveal three main findings: (1) speech identification is robust to acoustic interference, degrading only at very severe noise levels (i.e., negative SNRs); (2) the neural encoding of speech is enhanced for sounds carrying a clear phonetic identity compared to phonetically ambiguous tokens; and (3) categorical neural representations are more resistant to external noise than their non-categorical counterparts. Our findings suggest the mere process of categorization—a fundamental operation to all perceptual systems (Goldstone & Hendrickson, 2010)—aids figure-ground aspects of speech perception by fortifying abstract categories from the acoustic signal and making the speech code more resistant to external noise interference.

Behaviorally, we found listeners’ psychometric slopes were steeper when identifying clear compared to noise-degraded speech; identification functions became shallower only at the severe (negative) SNRs when noise levels exceeded that of speech. The resilience in perceptual identification suggests the strength of categorical representations is largely resistant to signal interference. Corroborating our modeling (Fig. 1), we found CP was affected only when the input signal was highly impoverished. These data converge with previous studies (Bidelman et al., under review; Gifford et al., 2014; Helie, 2017) suggesting category-level representations, which are by definition more abstract than their acoustic-sensory counterparts, are largely impervious to surface degradations. Indeed, as demonstrated recently in cochlear implant listeners, the sensory input can be highly impoverished, sparse in spectrotemporal detail, and intrinsically noisy (i.e., delivered electrically to the cochlea) yet still offer robust speech categorization (Han et al., 2016). Collectively, our data suggest that both the mere construction of perceptual objects and the natural discrete binning process of CP help category members to “pop out” amidst noise (e.g., Nothdurft, 1991; Perez-Gay et al., 2018) to maintain robust speech perception in noisy environments.

Noise-related decrements in CP (Fig. 3A) could reflect a weakening of internalized categories themselves (e.g., fuzzier match between signal and phonetic template) or alternatively, more general effects due to task complexity (e.g., increased cognitive load or listening effort; reduced vigilance). We can rule out the latter interpretation based on our RT data. The speed of listeners’ perceptual judgments to ambiguous speech tokens (Tk3) were nearly identical across conditions and invariant to noise (Fig. 3D). In contrast, RT functions became more categorical (“inverted V” pattern) with increasing SNR due entirely to changes in RTs for categorical members (continuum endpoints). These findings suggest that categories represent local *enhancements* of processing within the normal acoustic space (e.g., Fig. 1) which acts to sharpen categorical speech representations. That our data do not reflect gross changes in task vigilance is further supported by two additional findings: (i) lapses in performance did not vary across stimuli which suggests vigilance was maintained across conditions and (ii) ERPs predicted behavioral CP only for speech sounds that carried clear phonetic categories (Fig. 6). Therefore, the effects of noise on CP are most parsimoniously described as changes in the relative *sharpness* of the auditory categorical boundary (Livingston et al., 1998). That is, under extreme noise, speech identification is blurred, and the normal warping of the perceptual space is partially linearized, resulting in more continuous speech identification.

On the basis of fMRI, Guenther et al. (2004) posited that the length of time auditory cortical cells remain active after stimulus presentation might be shorter for category prototypes than for other sounds. They further speculated “the brain may be reducing the processing time for category prototypes, rather than reducing the number of cells representing the category prototypes (Guenther et al., 2004; p.55).” Some caution is warranted when interpreting these results given the sluggishness of the fMRI BOLD signal. Still, our data do not agree with Guenther et al. (2004)’s first assertion since ERPs showed larger (enhanced) activations to categorical prototypes within 200 ms. However, our RT data do agree with their second hypothesis. We found RTs were faster for prototypical speech (i.e., RT_Tk1/5_ < RT_Tk3_) providing confirmatory evidence that well-formed categories are processed more efficiently by the brain.

Our neuroimaging data revealed enhanced brain activity to phonetic (Tk1/5) relative to perceptually ambiguous (Tk3) speech tokens. This finding indicates categorical-level processing occurs as early as ∼150-200 ms after sound arrives at the ear (Alho et al., 2016; Bidelman et al., 2013; Toscano et al., 2018). Importantly, these results cannot be explained in terms of mere differences in exogenous stimulus properties. On the contrary, endpoint tokens of our continuum were actually the most distinct in terms of their acoustics. Yet, these prototypical stimuli elicited stronger neural activity than midpoint tokens (i.e., Tk1/5 > Tk3), which was not attributable to trivial differences in signal SNR. These results are broadly consistent with previous ERP studies (Altmann et al., 2014; Bidelman & Lee, 2015; Bidelman et al., 2013; Bidelman et al., 2014a; Bidelman et al., 2014b; Dehaene-Lambertz, 1997; Phillips et al., 2000), fMRI data (Binder et al., 2004; Kilian-Hütten et al., 2011), and near-field unit recordings (Bar-Yosef & Nelken, 2007; Chang et al., 2010; Micheyl et al., 2005; Steinschneider et al., 2003), which suggest auditory cortical responses code more than low-level acoustic features and reflect the early formation of auditory-perceptual objects and abstract sound categories.

ERP effects related to CP (Fig. 5) were consistent with activity arising from the primary and associative auditory cortices along the Sylvian fissure (Alain et al., 2017). The latency of these modulations was comparable to our previous electrophysiological studies on CP (Bidelman & Alain, 2015b; Bidelman et al., 2013; Bidelman & Walker, 2017) and may reflect a modulation of the P2 wave. P2 is associated with speech discrimination (Alain et al., 2010; Ben-David et al., 2011), sound object identification (Leung et al., 2013; Ross et al., 2013), and the earliest formation of categorical speech representations (Bidelman et al., 2013). Categorical neural enhancements (i.e., Tk1/5 > Tk3) also persisted ∼200 ms after P2, through what appeared to be a P3b-like deflection. Whether this wave reflects a late modulation of P2 or a true P3b response is unclear, the latter of which is typically evoked in oddball-type paradigms.

A similar “post-P2” wave (180-320 ms) has been observed during speech categorization tasks (Bidelman & Alain, 2015b; Bidelman et al., 2013), which varied with perceptual (rather) than acoustic classification. This response could represent integration or reconciliation of the input with a phonetic memory template (Bidelman & Alain, 2015b) and/or attentional reorienting during stimulus evaluation (Knight et al., 1989). Similar responses in this time window have also been reported during concurrent sound segregation tasks requiring active perceptual judgments of the number and quality of auditory objects (Alain et al., 2001; Alain et al., 2017; Bidelman & Alain, 2015a). The response might thus reflect controlled processes covering a widely distributed neural network including medial temporal lobe and superior temporal association cortices near parietal lobe (Alain et al., 2001; Dykstra et al., 2016). The posterior scalp distribution of this late deflection is consistent with this interpretation (Fig. 4). Paralleling the dynamics in our neural recordings, studies have shown that perceptual awareness of target signals embedded in noise produces early focal responses between 100-200 ms circumscribed to auditory cortex and posterolateral superior temporal gyrus that is followed by a broad, P3b-like response (starting ∼300 ms) associated with perceived targets (Dykstra et al., 2016). It has been suggested this later response, like the one observed here, is necessary to perceive target SIN or under the demands of higher perceptual load (Dykstra et al., 2016; Gutschalk & Dykstra, 2014; Lavie et al.).

What might be the mechanism for categorical neural enhancements (i.e., ERP_Tk1/5_ > ERP_Tk3_) and their high flexibility in noise? In their experiments on categorical learning, Livingston et al. (1998) suggested that when “category-relevant dimensions are not as distinctive, that is, when the boundary is particularly ‘noisy,’ a mechanism for enhancing separation may be more readily engaged” (p. 742). Phoneme category selectivity is observed early (<150 ms) (Alho et al., 2016; Bidelman et al., 2013; Chang et al., 2010), particularly in left inferior frontal gyrus (pars opercularis) (Alho et al., 2016), but only under active task engagement (Alho et al., 2016; Bidelman & Walker, 2017). While some nascent form of categorical-like processing may occur pre-attentively (Bizley & Cohen, 2013; Chang et al., 2010; Joanisse et al., 2007; Krishnan et al., 2009), it is clear that attention enhances the brain’s ability to form categories (Alho et al., 2016; Bidelman et al., 2013; Bidelman & Walker, 2017; Recanzone et al., 1993). In animal models, perceptual learning leads to an increase in the size of cortical representation and sharpening or tuning of auditory neurons for actively attended (but not passively trained) stimuli (Recanzone et al., 1993). We recently demonstrated visual cues from a talker’s face help sharpen sound categories to provide more robust speech identification in noisy environments (Bidelman et al., under review). While multisensory integration is one mechanism that can hone internalized speech representations to facilitate CP, our data here suggest that goal-directed attention is another.

The neural basis of CP likely depends on a strong audition-sensory memory interface (Bizley & Cohen, 2013; Chevillet et al., 2013; DeWitt & Rauschecker, 2012; Jiang et al., 2018) rather than cognitive faculties, *per se* (attentional switching and IQ; Kong & Edwards, 2016). Moreover, the degree to which listeners show categorical vs. gradient perception might reflect the strength of phonological processing, which could have ramifications for understanding certain clinical disorders that impair sound-to-meaning mapping (e.g., dyslexia; Calcus et al., 2016; Joanisse et al., 2000; Werker & Tees, 1987). CP deficits might be more prominent in noise (Calcus et al., 2016). Thus, while relations between CP and language-based learning disorders remains equivocal (Hakvoort et al., 2016; Noordenbos & Serniclaes, 2015), we speculate that assessing speech categorization under the taxing demands of noise might offer a more sensitive marker of impairment (e.g., Calcus et al., 2016).

More broadly, the noise-related effects observed here may account for other observations in the CP literature. For example, cross-language comparisons between native and non-native speakers’ CP demonstrate language-dependent enhancements in native listeners in the form of steeper behavioral identification functions (Bidelman & Lee, 2015; Iverson et al., 2003b; Xu et al., 2006) and more dichotomous (categorical) neural responses to native speech sounds (Bidelman & Lee, 2015; Zhang et al., 2011). Shallower categorical boundaries for non-native speakers can be parsimoniously described as changes in *intrinsic* noise, which mirror the effects of *extrinsic* noise in the current study. While the noise sources differ (exogenous vs. endogenous), both linearize the psychometric function and render speech identification more continuous. Similarly, the introduction of visual cues of a talker’s face can enhance speech categorization (Bidelman et al., under review; Massaro & Cohen, 1983). Such effects have been described as a reduction in decision ambiguity due to the mutual reinforcement of speech categories provided by concurrent phoneme-viseme information (Bidelman et al., under review). Future studies are needed to directly compare the impact of intrinsic vs. extrinsic noise on categorical speech processing. Still, the present study provides a linking hypothesis to test whether deficits (Calcus et al., 2016; Joanisse et al., 2000; Werker & Tees, 1987), experience-dependent plasticity (Bidelman & Lee, 2015; Xu et al., 2006), and effects of extrinsic acoustics on CP (present study) can be described via common physiological mechanisms.

## Acknowledgements

This work was supported by the National Institute on Deafness and Other Communication Disorders of the National Institutes of Health under award number NIH/NIDCD R01DC016267 (G.M.B.). We thank Gwyneth Lewis and Jared Carter for comments on earlier versions of this manuscript.

